# Advancing Variant Phenotyping in Myelodysplastic Syndromes via Computational Genomics of Mitochondrial Enzyme Complexes

**DOI:** 10.1101/2024.09.04.611279

**Authors:** Jing Dong, Michael T. Zimmermann, Neshatul Haque, Shahram Arsang-Jang, Wael Saber, Raul Urrutia

## Abstract

Mitochondria are essential cellular organelles that play critical roles in hematological disorders. Recurrent mutations in mitochondrial DNA (mtDNA) have been identified in patients with myelodysplastic syndromes (MDS) and serve as significant prognostic indicators for their outcomes following allogeneic hematopoietic stem-cell transplantation (allo-HCT). However, the biological mechanisms of mtDNA mutations remain unclear. The current study utilizes computational structural genomics to improve our understanding of pathogenic variants in mitochondria-encoded genes. This emerging genomics discipline employs structural models, molecular mechanic calculations, and accelerated molecular dynamic simulations to analyze gene products, focusing on their structures and motions that determine their function. We applied this methodology to perform deep variant phenotyping of entire mitochondria-encoded protein complexes associated with the pathobiology of MDS and their prognosis after HCT. Our results demonstrate that this approach significantly outperforms conventional analytical methods, providing enhanced and more accurate information to support the potential pathogenicity of these variants and better infer their dysfunctional mechanisms. We conclude that the adoption and further expansion of computational structural genomics approaches, as applied to the mitochondrial genome, have the potential to significantly increase our understanding of molecular mechanisms underlying the disease. Our study lays a foundation for translating mitochondrial biology into clinical applications, which is of significant mechanistic and biomedical relevance and should be considered in modern biomedical research.

---

Myelodysplastic syndromes (MDS) are a heterogeneous group of clonal hematopoietic cell disorders characterized by ineffective hematopoiesis, leading to blood cytopenias, and by a tendency to progress to acute myeloid leukemia (AML) in about 30% of patients.^1^ Despite advances in treatment, allogeneic hematopoietic stem-cell transplantation (allo-HCT) remains the only curative therapy for MDS. However, mortality after HCT is high due to disease relapse and transplant-related complications. Less than 50% of patients with MDS survive for five years after transplantation.^1^ Identifying which MDS patients will most likely benefit from allo-HCT is challenging, given the clinical and biological heterogeneity of the disease.^1^ Prognostic tools defining MDS risk groups, such as the International Prognostic Scoring System (IPSS) and the revised IPSS (IPSS-R), which have varying probabilities of survival and leukemic transformation are used to predict posttransplant outcomes for clinical decision-making. Such models rely on traditional clinical and hematologic parameters and, thus, cannot accurately predict outcomes. During the past decade, sequencing technologies have dramatically improved our understanding of the genetics underlying MDS, with more than 50 recurrently mutated genes being identified, most of which also associate with unfavorable outcomes after allo-HCT.^2^ Incorporating somatic mutations of 31 genes, Bernard *et al*. developed a clinical-molecular prognostic model [IPSS-Molecular (IPSS-M)], exhibiting improved prognostic discrimination across all clinical endpoints compared to the IPSS-R.^3^ Around the same time, the International Consensus Classification (ICC) published a revised framework for classifying myeloid neoplasms and acute leukemias, integrating genomic data.^4^ The shift to molecular markers may allow personalized medicine and has notably influenced clinical practice. Although it is now well recognized that genomic alterations play a central role in MDS, most studies have focused on characterizing mutations in the nuclear genome, overlooking the potential critical importance of the mitochondrial genome in achieving proper immunological competence and surveillance. Consequently, several laboratories, including ours, have developed an active program to better understand the structure, function, evolution, and pathogenicity that are associated with the mitochondrial genome in hematological diseases. Indeed, mitochondria are critical dynamic organelles maintaining hematopoietic cell homeostasis and differentiation.^5^ They produce cellular energy and reactive oxygen species *via* oxidative phosphorylation (OXPHOS), trigger apoptosis, and control heme biosynthesis and iron metabolism.^6^ Dysfunction in any of these processes is pathogenic for MDS.^6^ In addition, allo-HCT exacts a high metabolic demand to restore normal hematopoiesis and immunologic functions. Thus, mitochondria are not only the “powerhouse” of the cell, but also function in innate and adaptive immune response.^7^

Mitochondria possess their own DNA (mtDNA), a 16.6-kb circular double-stranded DNA encoding 22 transfer RNAs (tRNAs), two ribosomal RNAs (rRNAs), and 13 essential subunits for the mitochondrial electron transport chain (ETC). Alterations in mtDNA, including genetic mutations and epigenomic modifications, could impair mitochondrial function and contribute to disease pathophysiology and survival outcomes.^6,8^ For instance, we recently conducted whole-genome sequencing (WGS) to characterize the prognostic landscape of mitochondrial genome in MDS patients receiving allo-HCT and in their matched donors, which yielded insights into prognosis.^9^ Our findings demonstrate that genetic variants in frequently mutated mtDNA genes, such as mitochondrial Complex I genes (*e*.*g*., *MT-ND4L* and *MT-ND5*), are highly prognostic for allo-HCT outcomes in MDS patients.^9^ We further applied sequence-based (2D, two-dimensional) approaches (*e*.*g*., HmtVar, SIFT, PolyPhen-2, MutPred2, PANTHER, PhD-SNP and SNPs&GO) to predict the pathogenicity of the prognostic mtDNA variants, and identified 159 missense variants on mitochondrial Complex I genes that are potentially pathogenic.^9^ However, accurately predicting the mechanistic impact of these variants on molecular and cellular functions is even more challenging than predicting the effects of variants derived from the nuclear genome. The limitations of conventional annotation methods, which have historically been based primarily on sequence, do not allow for extensive evaluation of a mutated gene product in 3D (protein structure) or account for the molecular motions required for their function under normal conditions (molecular dynamics). This lack of molecular detail misses important context. With the advances in computational biophysics and AI-based fold prediction, as well as accelerated molecular dynamic simulations, our laboratories have demonstrated that approaches to genomics that account for these properties greatly enhance the functional interpretation of genomic variations.^10-13^

In our quest to advance the field of genomic interpretation, we have developed a multi-parametric mechanistic-based assessment of the structural and dynamic features of genetic variants to improve damaging predictions and gain insights into molecular dysfunction at the atomic level.^10-13^ We have successfully applied our 3D structural-based and 4D dynamics-based approach to reveal the molecular mechanisms of genetic variants in several nuclear genes (*e*.*g*., *KRAS, KDM6A, KMT2C*, and *RAG*) and shown that their performance is most often superior to 2D sequence-based techniques for identifying the underlying mechanisms of protein dysfunction.^10-13^ With the construction of the MT OXPHOS molecular machinery, the time is right for transformational advances in defining the molecular mechanisms of MT dysfunction. Here, we used our computational structural genomics (3D/4D) approaches to investigate the molecular mechanisms of mtDNA variants that were associated with posttransplant outcomes, with a specific focus on mitochondrial genes that form the OXPHOS Complex I, the largest component of the mitochondrial OXPHOS system.

In this study, we demonstrate the analytical power of this methodology as applied to mitochondria genomics. For this purpose, the Cryo-EM structure of mitochondrial respiratory supercomplex, composed of three membrane-bound members of the ETC (*i*.*e*., complex I, dimeric complex III, and complex IV), were retrieved from RCSB Protein Data Bank (PDB; entry code: 5xth)^14^ and used in the current analysis to give the most physiologically relevant context to our 3D assessment and structure-based calculations (**Figure 1A**). For each mitochondrial complex I gene (*i*.*e*., *MT-ND1, MT-ND2, MT-ND3, MT-ND4, MT-ND4L, MT-ND5*, and *MT-ND6*), we selected one potentially pathogenic missense variant from our previous study^9^ that has the highest deleterious predicted score according to 2D annotation methods (*i*.*e*., *MT-ND1* m.3470T>C, p.L55P; *MT-ND2* m.5016T>C, p.S183P; *MT-ND3* m.10176G>A, p.G40S; *MT-ND4* m.11423G>A, p.E222K; *MT-ND4L* m.10726G>A, p.G86D; *MT-ND5* m.12770A>G, p.E145G; and *MT-ND6* m.14601G>A, p.P25S). Selected variants entered 3D structure-based assessment, as a proof of concept.

**Figure 1.**
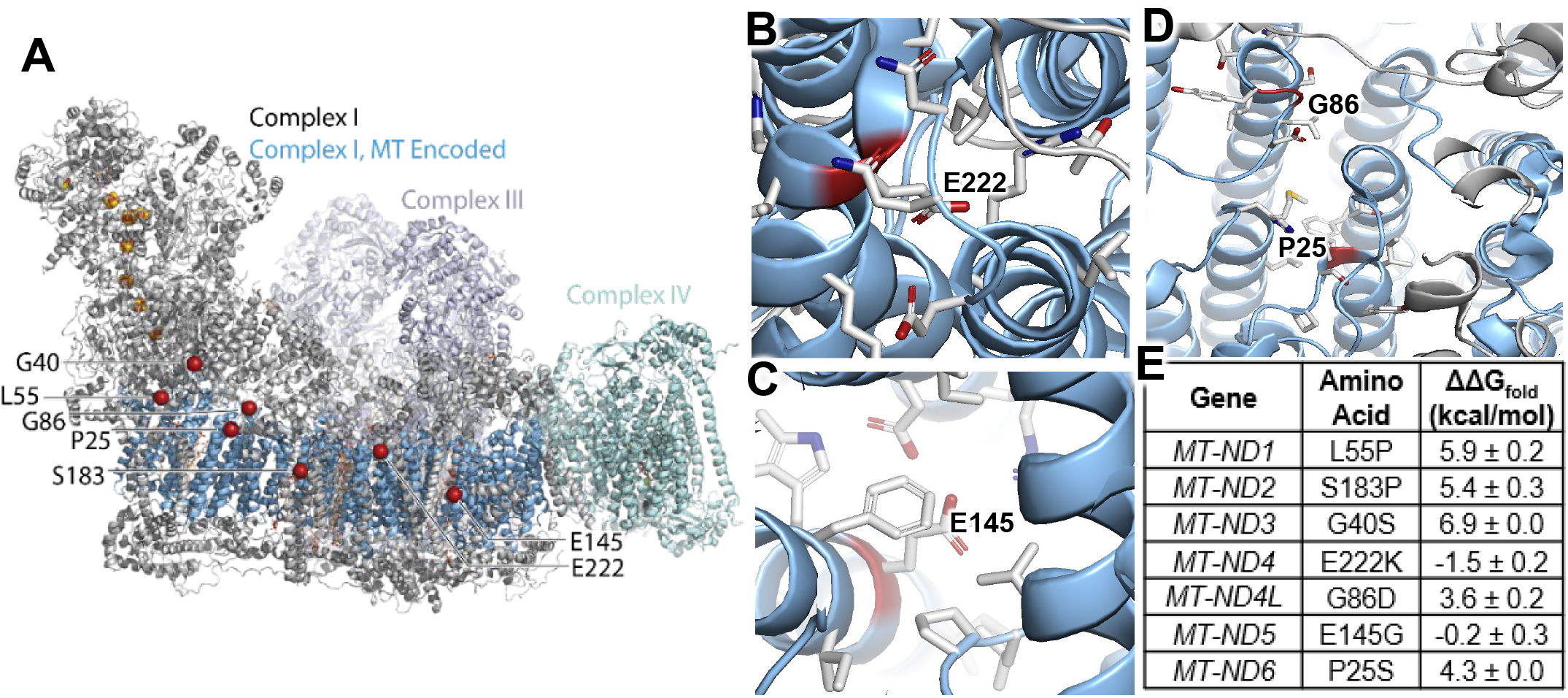
Structure of the MT super-complex reveals that MDS VUS are distributed throughout its quaternary structure. **A)** We label each selected variant with a red sphere marking its location. **B)** MT-ND4:E222 and **C)** MT-ND5:E145 are located within internal pockets that are lined with combinations of positive and negative charges, likely facilitating the flexible conformational changes that are required for electron transport. **D)** Selected mutations are nearby in space and typically at the ends of alpha helices where they likely modulate flexibility. **E)** Structure-based stability calculations reveal the disruption to the complex by MDS-associated variation.

To investigate the potential damage caused by these mtDNA mutations on their protein structures, we used an integrative approach to calculate the multifaceted effects that a genetic change can have on its encoded product. We used well-established metrics from protein structural analysis (*i*.*e*., amino acid local packing density, solvation energy, hydrogen bonds, and salt bridges) and more advanced metrics (*i*.*e*., multi-body statistical contact potentials, energetic frustration, and thermodynamic cooperativity) and made images using PyMOL v2.5.4.^15-20^ By examining the environments around the selected mtDNA variants, we found that mutations of *MT-ND4*:E222 (**Figure 1B**) and *MT-ND5*:E145 (**Figure 1C**) were likely to change the dynamics of the complex because they are internally-facing charges surrounded by a mixture of positive and negative charges. The residue E222 is in a solvent exposed helix and appears to have a crucial role in holding the adjacent solvent exposed helix by hydrogen bond. Additionally, the residue E145 is located at the interface stabilizing the interaction with MT-ND4. We anticipate that such an arrangement balances architectural stability with essential flexibility. Mutation to a different charge (*e*.*g*., E222K) or that form greater internal cavities (*e*.*g*., E145G) will have dynamic effects on the structure or the electron transport process. Interestingly, these candidate variants were selected without structural data, and there are proximal relationships among them like *MT-ND6*:P25 being across from *MT-ND4L*:G86 (**Figure 1D**). The role of G86 is probably to provide a steric free edge at the end of helix for a sharp turn of C-terminal peptide, which could otherwise not be stabilized if placed in other direction. We anticipate that any bulky mutation at this site could be deleterious. Additionally, we performed evolutionary coupling analysis (ECA) with tools from the Marks lab,^21^ allosteric path analysis with tools from the Bahar lab,^22,23^ and calculated structural stability using the same tools plus those of Rousseau.^24,25^ These ECA identified all seven variants as deleterious. Moreover, allosteric path analysis identified six out of seven variants as deleterious with one neutral call (*MT-ND4*:E222K) but within 1% of the threshold required for assignment to this category. Structural stability analysis, which is based on molecular mechanic calculations of folding energy identified one variant with neutral (*MT-ND5*:E145G), one stabilizing (*MT-ND4*:E222K) and the remaining five variants with highly destabilizing effect (**Figure 1E**). These data indicate that the prognostic effects of these mtDNA variants on MDS patients receiving allo-HCT likely involve their deleterious impacts on mitochondrial function. Thus, when using computational structural genomics for their analyses, mtDNA variants are a powerful prognostic tool for predicting the outcome of allo-HCT and advances the field of precision medicine for MDS.

Considering the novelty of these results, it becomes important to briefly highlight additional applications. Computational structural genomics increases our ability to better predict pathogenicity than previous methods due to the precision of these methods over the ones popularized by guidelines. More importantly, computational structural genomics provides explanations for mechanisms of dysfunction underlying the impact of variants, in particular because of its used of advanced methods of molecular dynamic simulations. Lastly, since many drugs have been developed using similar methodologies, the conformations that results from molecular dynamics simulation build the trajectory toward the identification of mutant-specific cavities for development of organic probes or small drugs. Therefore, we anticipate that the continue application to the deep variant characterization of both nuclear and Mt-DNA-encoded mitochondria proteins will significantly advance not only the field of MDS but other genomic diseases of importance to our field.

In conclusion, in the current molecular era, mtDNA variants are a powerful prognostic tool for both predicting the outcome of allo-HCT and represent an advancing the field of precision medicine for MDS. Studies such as we illustrated here increase our ability to better predict pathogenicity, provide explanations for their mechanism of dysfunction, and yield structural conformation to help build novel therapeutic tools. In this manner, we can better leverage mtDNA genomics into the functional interpretation and actionable potential. Thus, we are optimistic that this study will serve as a useful framework for characterizing the spectrum of translational research, from bedside (*e*.*g*., measuring individual’s genomics) to bench (*e*.*g*., computational structural genomics) then back to bedside (*e*.*g*., improved risk stratification and clinical outcomes), necessary for our precision medicine practice and research. Such work requires a multidisciplinary team with high level of expertise in HCT clinical management, molecular epidemiology, systems biology, mitochondrial biochemistry, computational biophysics, computational chemistry, and stem cell biology. Should such a proposed translational cycle be successful, it will be a paradigm shift in allo-HCT, from donor selection, recipient risk stratification to mitochondrial targeted therapying in HCT, MDS, and most likely to other diseases caused by alterations in this organelle.

## Competing interests

None

## Authors’ Contributions

J.D., M.T.Z. and R.U. conceived the study and oversaw the project; M.T.Z., N.H. and R.U. designed the computational framework and analyze the data; J.D., M.T.Z., N.H., S.A., W.S, and R.U. wrote and revised and manuscript; All authors read and approved the final manuscript.

## Acknowledgements

Jing Dong is supported by the NHLBI (K01 HL164972) and the Medical College of Wisconsin Cancer Center. This project was funded in part by the National Institutes of Health Grant R01 DK52913, and the Linda T. and John A. Mellowes Center for Genomic Sciences and Precision Medicine at the Medical College of Wisconsin.

## Notes

### Competing Interest Statement

The authors have declared no competing interest.

